# Small molecule FGF23 inhibitors increase serum phosphate and improve skeletal abnormalities in *Hyp* mice

**DOI:** 10.1101/2020.08.04.236877

**Authors:** Zhousheng Xiao, Jiawang Liu, Shih-Hsien Liu, Loukas Petridis, Chun Cai, Li Cao, Guangwei Wang, Ai Lin Chin, Jacob W. Cleveland, Munachi O. Ikedionwu, Jesse D. Carrick, Jeremy C. Smith, L. Darryl Quarles

**Affiliations:** Department of Medicine, College of Medicine, University of Tennessee Health Science Center, Memphis, TN, 38165, USA; Department of Pharmaceutical Sciences, College of Pharmacy, University of Tennessee Health Science Center, Memphis, TN, 38163, USA; UT/ORNL Center for Molecular Biophysics, Oak Ridge National Laboratory, 1 Bethel Valley Road, Oak Ridge, Tennessee 37830, USA; Department of Biochemistry and Cellular and Molecular Biology, University of Tennessee, Knoxville TN 37996, U.S.A.; Department of Chemistry, Tennessee Technological University, 55 University Drive, Cookeville, Tennessee 38501, USA

**Keywords:** FGF23, FGF23 Inhibitor, Phosphate Homeostasis, Hypophosphatemic Rickets

## Abstract

Fibroblast growth factor 23 (FGF23) is a bone-derived hormone that binds to binary FGFR/α-Klotho receptor complexes in the kidney tubules to inhibit phosphate reabsorption and 1,25(OH)_2_D production. Excess FGF23 causes X-linked hypophosphatemia (XLH) and tumor induced osteomalacia (TIO). Recently, Burosumab, an FGF23 blocking antibody, was approved for treating these hypophosphatemic disorders. A small molecule inhibitor that specifically binds to FGF23 to prevent activation of FGFR/α-Klotho complexes has potential advantages over a systemically administered FGF23 blocking antibody. We previously identified the small molecule ZINC13407541 (N-[[2-(2-phenylethenyl)cyclopenten-1-yl]methylidene]hydroxylamine) as a therapeutic lead FGF23 antagonist. Additional structure-activity studies developed a series of ZINC13407541 analogues with enhanced drug-like properties. In this study, we tested in a pre-clinical *Hyp* mouse homologue of XLH a direct connect analogue (**8n**) [(*E*)-2-(4-(*tert*-butyl)phenyl)cyclopent-1-ene-1-carbaldehyde oxime] that exhibited the greatest stability in microsomal assays, and **13a** [(*E*)-2-((*E*)-4-methylstyryl)benzaldehyde oxime] that exhibited increased *in vitro* potency. We found that pharmacological inhibition of FGF23 with either of these compounds blocked FGF23 signaling and significantly increased serum phosphate and 1,25(OH)_2_D concentrations in *Hyp* mice. Long-term parenteral treatment with **8n** or **13a** also enhanced linear bone growth, increased mineralization of bone, and narrowed the growth plate in *Hyp* mice. The more potent **13a** compound showed greater therapeutic efficacy in *Hyp* mice. Further optimization of these FGF23 inhibitors may lead to versatile drugs to treat FGF23-mediated disorders.

## INTRODUCTION

Fibroblast Growth Factor 23 (FGF23) is a hormone produced by osteoblasts/osteocytes in the bone that activates FGF receptor/α-Klotho (Kl) binary complexes in renal proximal tubules to regulate phosphate reabsorption and 1,25(OH)_2_D metabolism and in distal tubules to adjust sodium and calcium reabsorption ^(1, 2)^. FGF23 plays a causal role in hereditary hypophosphatemic disorders, such as X-linked (XLH) and autosomal recessive (ARH) hypophosphatemic rickets and acquired tumor-induced osteomalacia (TIO). Elevated FGF23 plays an adaptive role in maintaining phosphate homeostasis in chronic kidney disease and is associated with left ventricular hypertrophy and increased cardiovascular mortality in this setting ^(3–7)^.

Until recently, treatment of these hypophosphatemic disorders consisted of 1,25(OH)_2_D and phosphate supplements. This approach does not cure the disease and is associated with toxicities related to excess phosphate and 1,25(OH)_2_D, including nephrocalcinosis ^(8, 9)^. TIO can be cured by resection of the tumor that is producing FGF23, but surgical removal of the tumor is not possible in approximately 50% of these patients ^(10, 11)^.

Recent therapeutic advances use FGF23 blocking antibodies to treat disorders of FGF23 excess. An FGF23 blocking antibody KRN23 (Burosumab) has been recently approved for the treatment of XLH and TIO by the FDA^(12, 13)^. KRN23 subcutaneously administered at an average dose of 0.98 mg/kg every two weeks improves rickets and increases serum phosphate levels in XLH ^(14)^, and was superior to conventional treatment with phosphate and 1,25(OH)_2_D supplements. To date, KRN23 has not been associated with toxicity at the doses used to treat hypophosphatemia, but the initial use of a high affinity FGF23 blocking antibody in pre-clinical rodent models resulted in excess mortality ^(15)^, principally due to over-suppression of FGF23 resulting in hyperphosphatemia and 1,25(OH)_2_D toxicity ^(16–18)^. Additional disadvantages of KRN23 include the need for systemic administration, long-half life (elimination half-life of 18 days), and cost.

Developing an orally, bioavailable small molecule drug to block FGF23 would have several potential advantages over KRN23. To this end, using homology modeling, molecular dynamics simulation, and virtual high-throughput screening we identified ZINC13407541 (N-[[2-(2-phenylethenyl)cyclopenten-1-yl]methylidene]hydroxylamine) which was experimentally validated to bind to FGF23 and inhibit its interaction with FGFRs ^(19)^. Subsequent medicinal chemistry investigation of structure-activity relationships based on ZINC13407541 produced 36 analogues with different stabilities and potencies. The core ZINC13407541 structure contains two ring systems, bridged with an ethylene, and a carbaldehyde oxime functional group on the *ortho*-position of ring A that is essential for activity (**Figure 1**). Replacing a five-membered aliphatic ring A with a six-membered ring or an aromatic system markedly increases potency of the FGF23 antagonists. A substitution of the electron-donating groups (CH3 or OCH3) on para-position of aryl ring B increased the *in vitro* FGF23 inhibitor effects. Removal of the ethylene bridge to create a direct connect derivative decreased efficacy, but increased metabolic stability ^(20)^.

**Figure 1.**
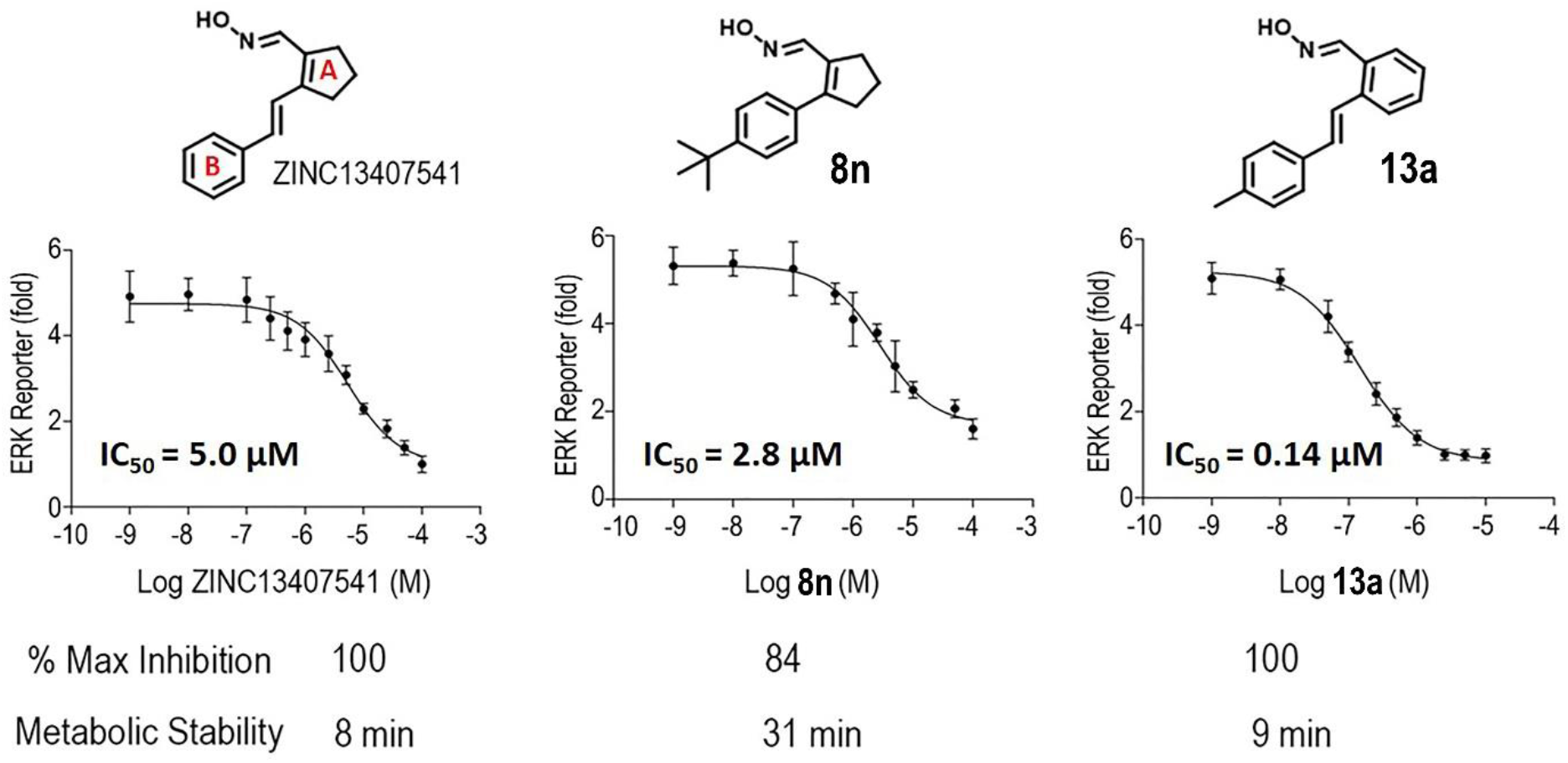
Structure, *in vitro* efficacy, and metabolic stability of ZINC13407541 and its analogues **8n** and **13a**.

In this study, we tested the *in vivo* efficacy of the direct connect analogue (**8n)** [(*E*)-2-(4-(*tert*-butyl)phenyl)cyclopent-1-ene-1-carbaldehyde oxime], eliminating the ethylene bridge between the two ring systems of ZINC13407541, and the most potent compound (**13a)** [(*E*)-2-((*E*)-4-methylstyryl)benzaldehyde oxime] in the preclinical *Hyp* mouse model of XLH.

## RESULTS

### Pharmacokinetics, *in vitro* metabolic stability and IC_50_

We performed murine pharmacokinetics of the lead compound ZINC13407541. ZINC13407541 was administered as a single intravenous dose (1 mg/kg) and plasma concentrations were determined by LC-MS/MS. ZINC13407541 was found to have a half-life of ≈2 hours (**Figure 1S**).

Structure activity relationship (SAR) studies generated 36 compounds with different stability and potency in the *in vitro* assays ^(20)^. Compound (**8n**) [(*E*)-2-(4-(*tert*-butyl)phenyl)cyclopent-1-ene-1-carbaldehyde oxime], a direct connect analogue of ZINC13407541showed the greatest stability and **13a** [(*E*)-2-((*E*)-4-methylstyryl)benzaldehyde oxime] the greatest potency of the 36 compounds generated.

As shown in **Figure 1**, compound **8n** has eliminated the ethylene bridge to form a directly connected two-ring scaffold, which decreased maximum inhibition activity (% Maximum inhibition 84%), but showed increased IC_50_ (2.8 μM) and metabolic stability (31 minutes) compared to ZINC13407541 (100%, 5.0 μM, and 8 min). In contrast, compound **13a** has exchanged the cyclopentene with a phenyl group, showing no changes in metabolic stability (9 minutes) and similar maximum inhibition activity (% Maximum inhibition 100%), but 36-fold higher potency for inhibiting FGF23 activity (**13a**, IC_50_= 0.14 μM) compared with ZINC13407541 (IC_50_ = 5.0 μM).

### Molecular docking of experimentally verified hits on the N-terminal domain of FGF23

Analysis of our structural model of FGF23 shows that Gln156 is a consensus binding site on the N-terminal domain of FGF23 for ZINC13407541 and its two analogs, **13a** and **8n** (**Figure 2**). ZINC13407541 and **13a** adopt similar conformations *in silico*. Further, ZINC13407541 (**Figure 2B**) and **13a** (**Figure 2C**) each have two hydrogen bonds interactions with Gln156 and Asn122, and compound **8n** (**Figure 2D**) has one hydrogen bond interaction with Gln156. **Table 1** shows the estimated free energy of binding (ΔG) of the three compounds on FGF23 using two different methods: AutoDock VinaMPI ^(21, 22)^ and K_DEEP_ ^(23)^. The differences in ΔG between the compounds are within the uncertainties associated with the methods used ^(22,23)^. Taken together, our computational results show that these three compounds are likely to bind to Gln156 on FGF23 with fairly similar binding affinities.

**Figure 2.**
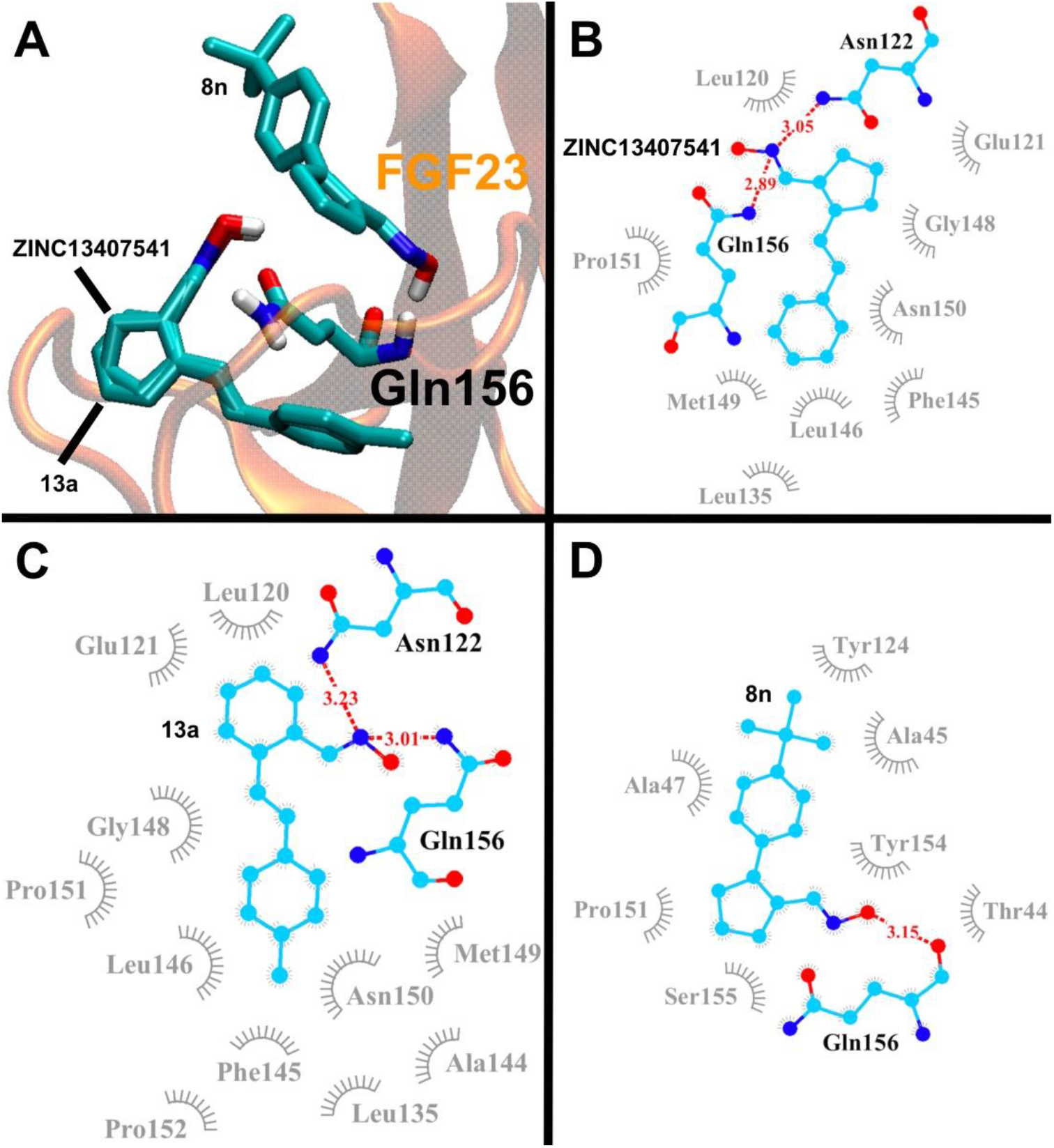
The computationally predicted interaction of ZINC13407541 and its two analogs, **13a** and **8n**, with the N-terminal domain of FGF23 (PDB code: 5W21) shown in **A**: 3D structure; in **B**, **C**, and **D**: 2D residue-contacting map for ZINC13407541, **13a**, and **8n**, respectively. Hydrogen bonds are shown in red dash lines with donor-acceptor distances in Å. Hydrophobic interactions are shown in gray.

**Table 1.**
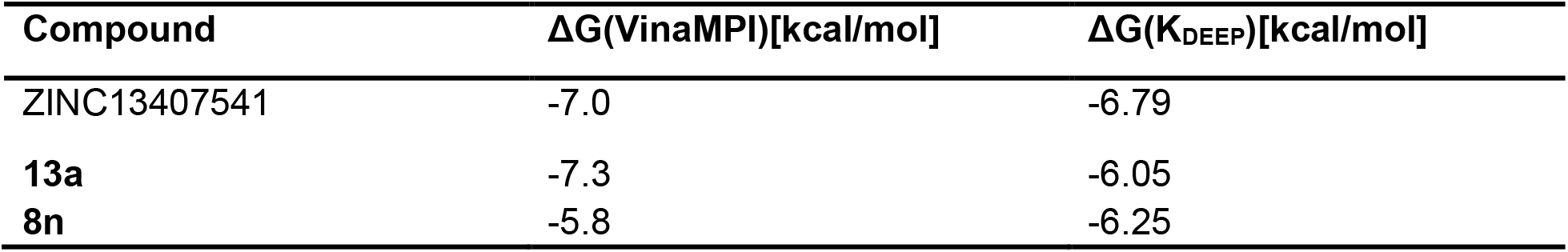
The computationally estimated free energy of binding (ΔG) of ZINC13407541, 13a, and 8n in the N-terminal dpmain of FGF23 using AutoDock Vina MPI and K_DEEP_. The corresponding structures are shown in Figure 2.

### Mineral ion homeostasis changes in serum of *Hyp* mice treated with single-dose or short-term FGF23 inhibitors

Next, we examined the *in vivo* efficacy of both **8n** and **13a** analogues compared to the parent molecule ZINC13407541 in preclinical animal studies. In *Hyp* mice a single intraperitoneal (IP) injection of ZINC13407541 resulted in increasing serum phosphate levels 2 hours after administration, peaked 4 hours after treatment, and returned to original levels by 24 hours post injection (**Figure 3A and 3B**). Also, the single IP injection of ZINC13407541 increased *Hyp* mice serum phosphate levels in a dose-dependent manner, the maximum effect achieved in the dose of 100 mg/kg ZINC13407541 (**Figure 3A and 3B**). In contrast, the ZINC13407541 treatment did not affect serum calcium levels (**Figure 3C and 3D**). Unexpectedly, *Hyp* mice with 100 mg/kg dosing of ZINC13407541 after 24 hours or 200 mg/kg dosing of ZINC13407541 after 4 hours exhibited a 2-fold increase in serum FGF23 levels (**Figure 3E and 3F**). Compound **13a** showed same time- and dose-dependent responses similar to what we observed with ZINC13407541, but had a greater magnitude in the increase in serum phosphate levels in *Hyp* mice 4 hours after dosing, consistent with its improved IC_50_ (**Figure 4A and 4B**). At 100 mg/kg **13a** almost completely corrected the phosphate levels of *Hyp* mice. No changes were observed in serum calcium and FGF23 levels in *Hyp* mice after administration (**Figure 4C-4F**). Subsequently, we performed short-term treatments with ZINC13407541 (100 mg/kg), **8n** (100 mg/kg), or **13a** (100 mg/kg) IP injection twice a day for 3 days. We obtained similar effects of these three compounds in increasing serum phosphate levels in *Hyp* mice (**Figure 5A and 5B**). No changes in serum calcium or FGF23 levels were observed with this short-term treatment regimen in *Hyp* mice (**Figure 5C-5F**). In long-term exposure studies, we decided to use half the effective dose (50 mg/kg) of ZINC13407541, **8n**, and **13a** IP injection twice a day for 4-weeks in *Hyp* mice to assess potential skeletal effects.

**Figure 3.**
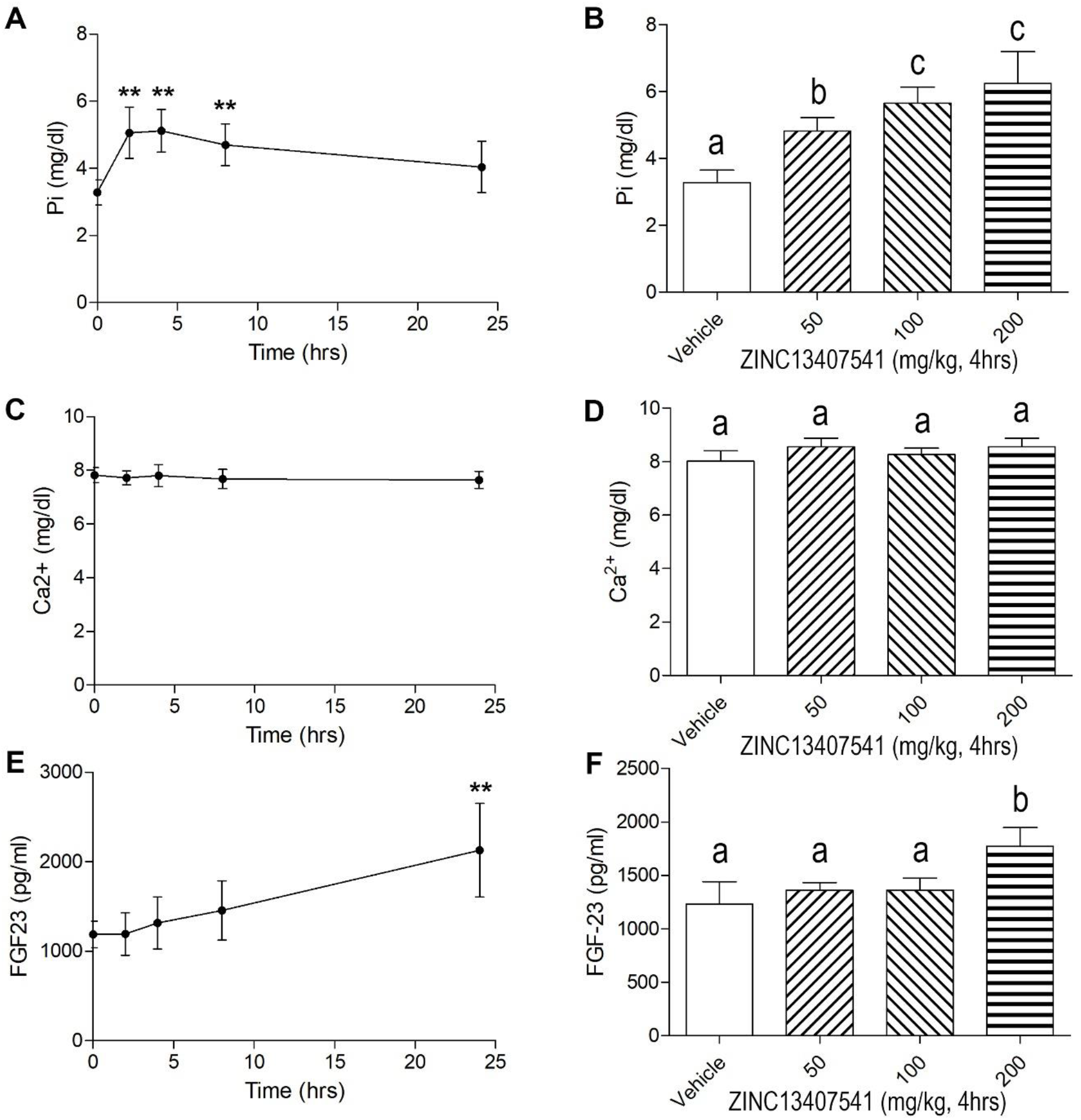
Time and dose dependent effects of ZINC13407541 on mineral ion homeostasis and FGF23 levels in *Hyp* mice. *, **, *** indicates significant difference from vehicle control group. Values (mean ± SD, n=5-6) with the different superscript are significantly different at *P*<0.05.

**Figure 4.**
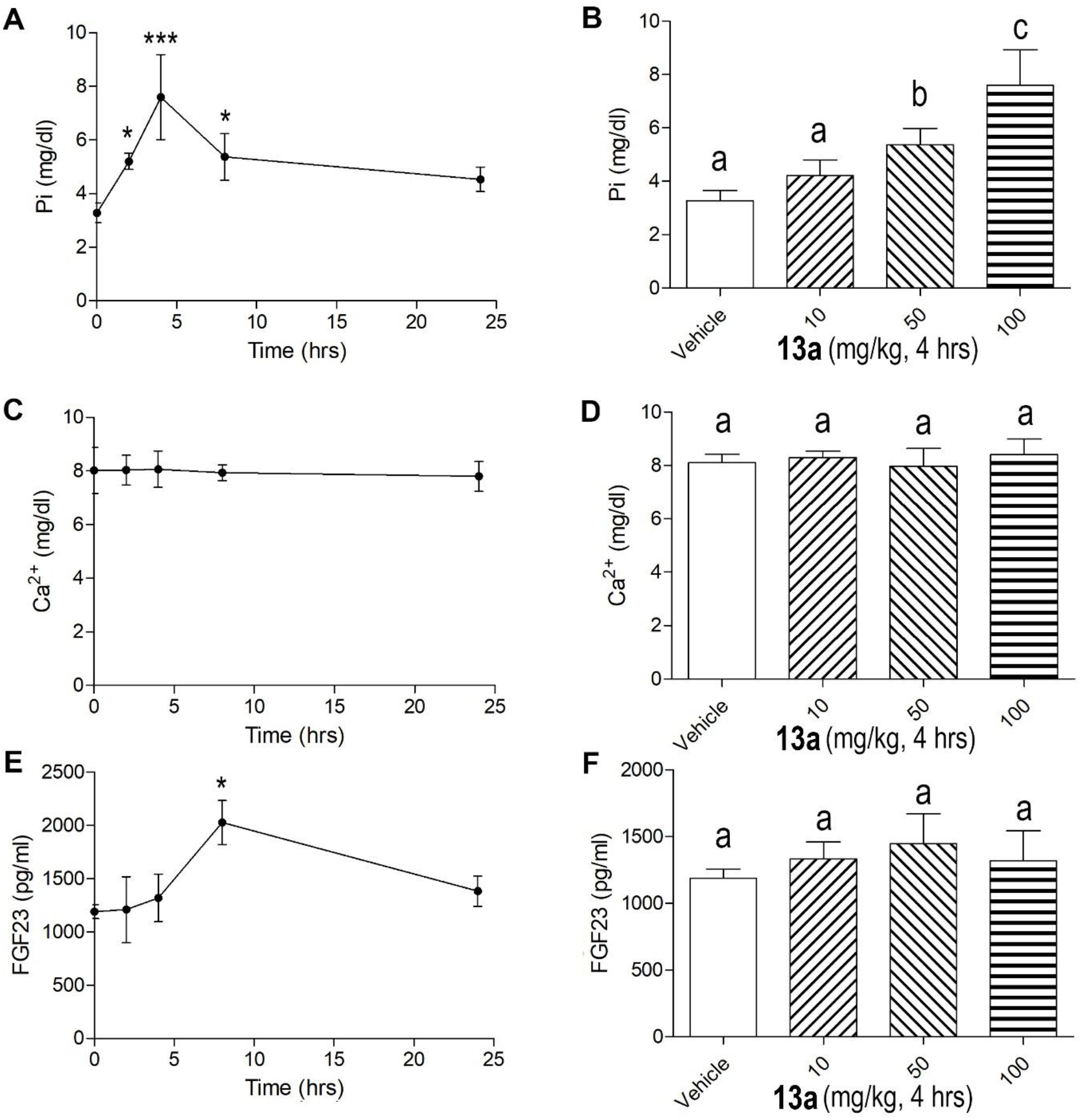
Time and dose dependent effects of **13a** on mineral ion homeostasis and FGF23 levels in *Hyp* mice. *, **, *** indicates significant difference from vehicle control group. Values (mean ± SD, n=5-6) with the different superscript are significantly different at *P*<0.05.

**Figure 5.**
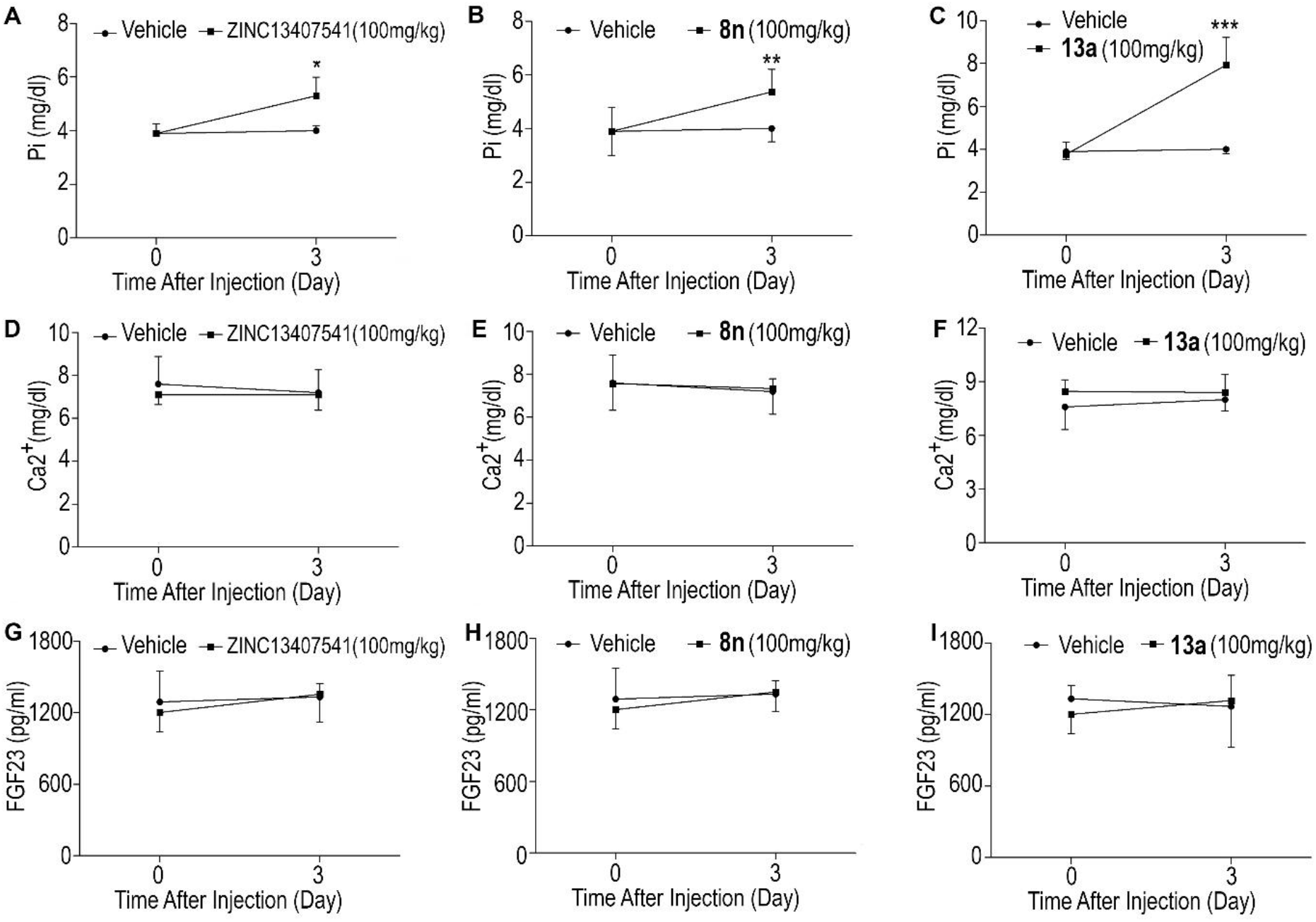
Short-term effects of ZINC13407541, **8n**, and **13a** on mineral ion homeostasis and FGF23 levels in *Hyp* mice. Data are expressed as the mean ± S.D. from 5-6 mice. *, **, *** indicates significant difference from vehicle control group.

### Serum biochemistry and skeletal phenotype changes of *Hyp* mice treated long-term with FGF23 inhibitors

We treated 4- to 8-week-old *Hyp* mice with either vehicle control, ZINC13407541 (50mg/kg), **8n** (50mg/kg), or **13a** (50mg/kg) for 4 weeks followed by measurement of serum biochemistry and assessment of skeletal features. Serum phosphate levels were significantly elevated in all treated groups compared to controls. The increase in serum phosphate was significantly greater in **13a** compared to ZINC13407541 and **8n** treated mice. No significant changes of FGF23 levels were observed in **8n** and **13a** treated groups compared to vehicle treated mice (**Table 2**). There was a 1.5-fold increase of FGF23 in the ZINC13407541 treated group, similar to previous observations ^(24)^. FGF23 is known to suppress parathyroid hormone (PTH) and active form (1,25(OH)2D) of vitamin D levels. Consistent with inhibition of FGF23 signaling, both PTH and 1,25(OH)_2_D levels were markedly increased in ZINC13407541, **8n** and **13a** treated mice. FGF23 also has distal tubular effects, increasing sodium reabsorption and suppressing aldosterone levels ^(25, 26)^. ZINC13407541, **8n**, and **13a** resulted in significant stimulation in aldosterone levels compared to vehicle treated mice (**Table 2**).

**Table 2.** Serum biochemistry of *Hyp* mice treated with ZINC13407541 & its analogues.

| Parameters | Vehicle | ZINC13407541 | 8n | 13a | *p*-value |
| --- | --- | --- | --- | --- | --- |
| FGF23 (pg/ml) | 1413±366<sup>a</sup> | 2359±861<sup>b</sup> | 1754±445<sup>a</sup> | 1378±289<sup>a</sup> | 0.0110 |
| Phosphate (mg/dl) | 4.6±0.60<sup>a</sup> | 6.5±1.02<sup>b</sup> | 6.7±0.58<sup>b</sup> | 7.8±1.08<sup>c</sup> | <0.0001 |
| Calcium (mg/dl) | 8.0±1.23<sup>a</sup> | 8.9±1.22<sup>a</sup> | 8.4±1.77<sup>a</sup> | 8.3±1.63<sup>a</sup> | 0.7208 |
| PTH (pg/ml) | 78±12<sup>a</sup> | 122±21<sup>b</sup> | 118±20<sup>b</sup> | 138±21<sup>b</sup> | 0.0002 |
| 1,25(OH)<sub>2</sub>D (pg/ml) | 19±13<sup>a</sup> | 65±14<sup>b</sup> | 69±19<sup>b</sup> | 116±14<sup>c</sup> | <0.0001 |
| Aldosterone (pg/ml) | 278±84<sup>a</sup> | 389±68<sup>b</sup> | 401±113<sup>b</sup> | 497±71<sup>b</sup> | 0.0031 |

Data are mean ±S.D. from 5-6 serum samples of the ZINC13407541 & its analogs treated individual mice. Values with the different superscript are significantly different at *P*<0.05.

With regards to skeletal effects, we observed that body weight, body length, femur length, and tail length were significantly increased in ZINC13407541, **8n**, or **13a** treated groups compared to vehicle controls at the end of the treatment period (**Figure 6**). *Hyp* mice treated with ZINC13407541, **8n**, or **13a** showed 15%, 18%, and 30% increments in femoral bone mineral density (BMD), respectively, compared to vehicle controls (**Figure 7A**). Micro-CT 3D images revealed that both ZINC13407541 and **8n** treated groups had similar increases in trabecular bone volume (TBV, 51%) and cortical bone thickness (Ct.Th, 30%). The *Hyp* mice treated with **13a** had greater increases in both TBV (78%) and Ct.Th (44%) than either ZINC13407541 or **8n** group (**Figure 7B**). The width of the growth plate was significantly decreased following treatment with ZINC13407541, **8n**, or **13a**. In summary, **13a** had greater therapeutic effects on the healing of the growth plate in *Hyp* mice (**Figure 7B**).

**Figure 6.**
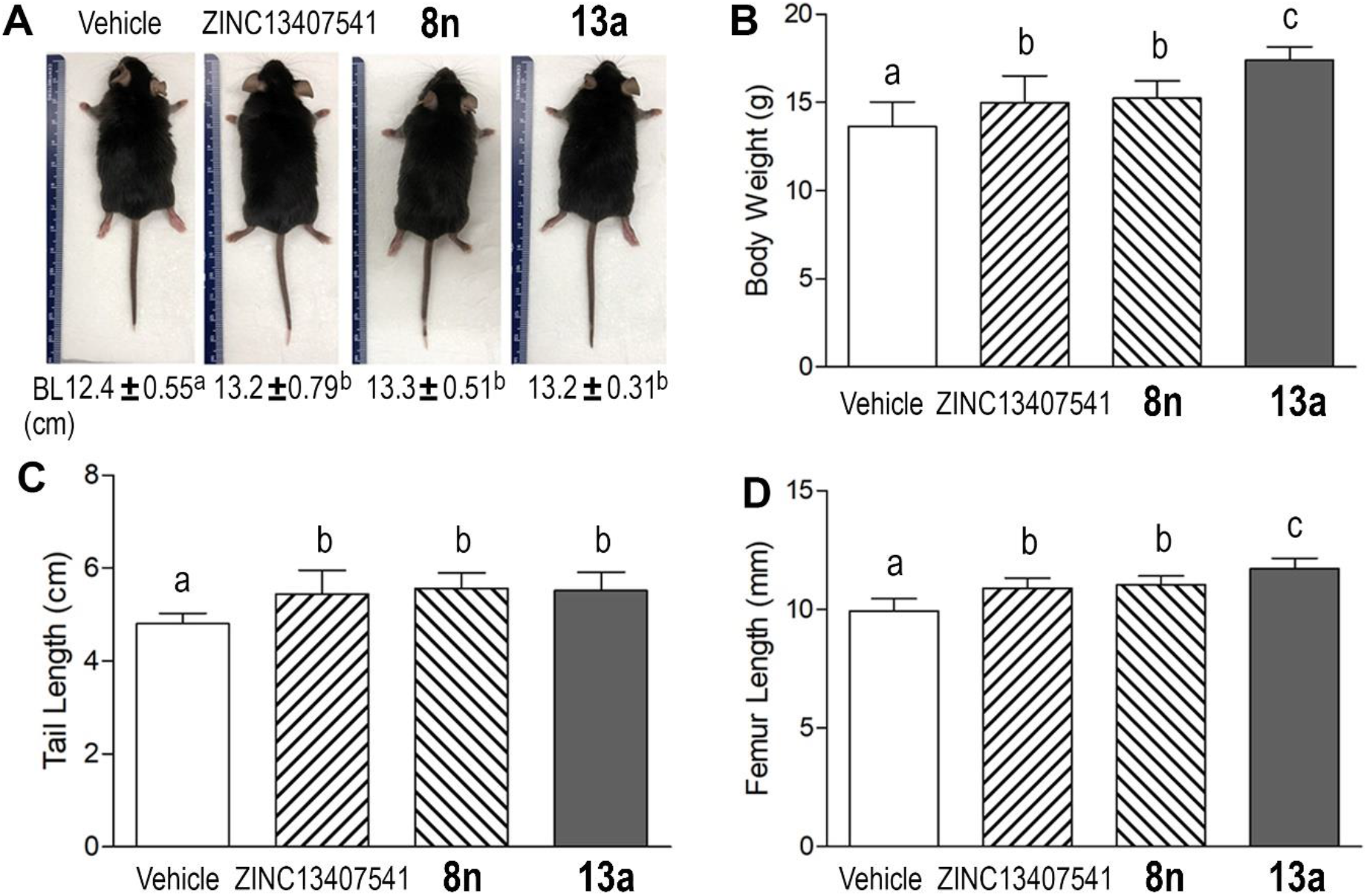
Long-term effects of ZINC13407541, **8n**, and **13a** on body length (BL) (A), body weight (B), tail length (C), and femur length (D) in *Hyp* mice. Data are expressed as the mean ± S.D. from 6-10 mice. Values with the different superscript are significantly different at *P*<0.05.

**Figure 7.**
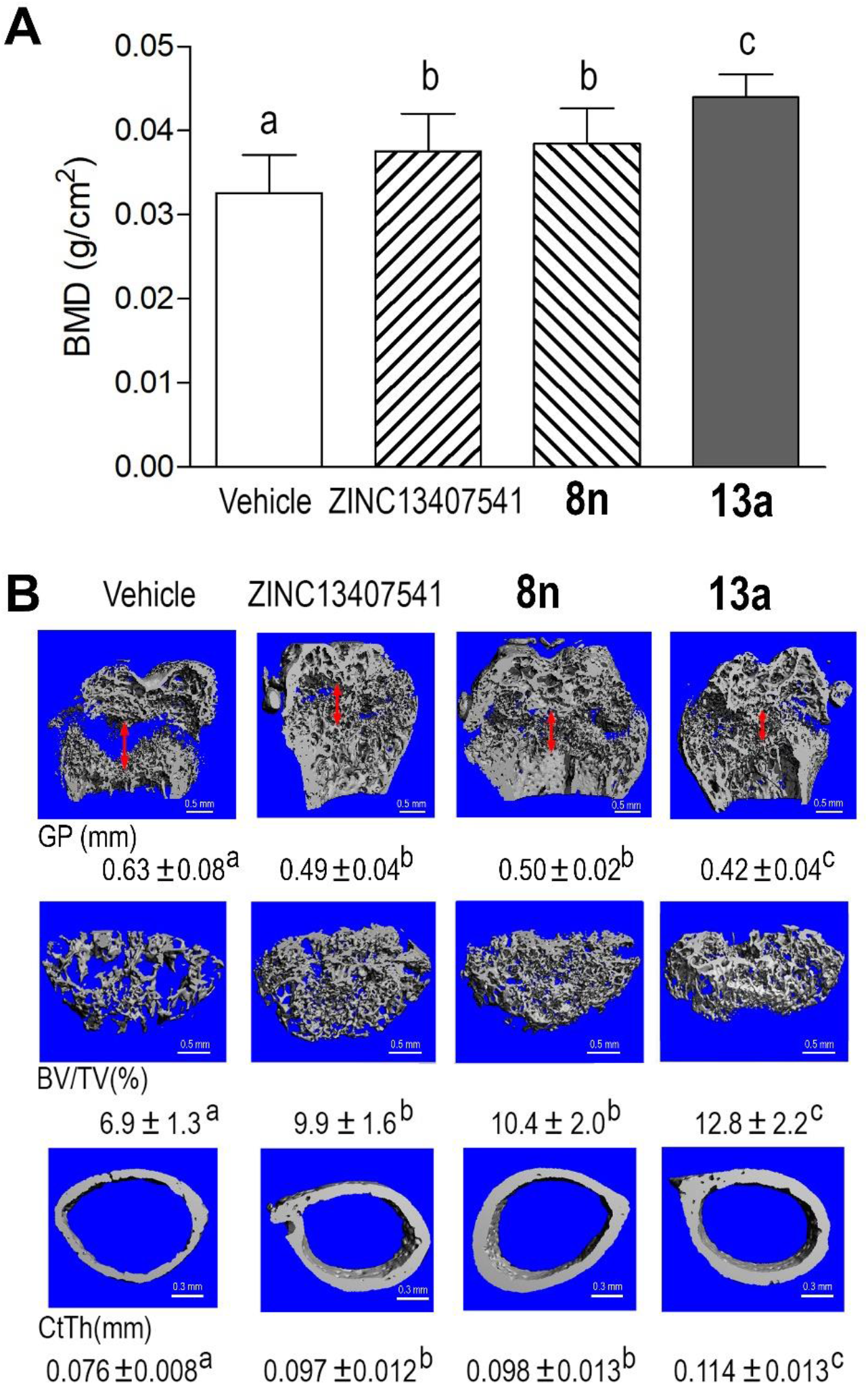
Long-term treatment of ZINC13407541, **8n**, and **13a** on bone mineral density (BMD), width of the growth plate (GP, double red arrow), femoral bone volume (BV/TV), and cortical thickness (CtTh) in *Hyp* mice. (A) BMD, (B) Micro-CT 3D images. Data are expressed as the mean ± S.D. from 6-10 mice. Values with the different superscript are significantly different at *P*<0.05.

### Gene expression profiles changes in bone of *Hyp* mice treated long-term with FGF23 inhibitors

*Hyp* mice are known to have increased FGFR1 signaling and Wnt/β-catenin signaling, and a characteristic gene expression profile in bone associated with the presence of rickets/ osteomalacia ^(27–29)^. Real-time RT-PCR revealed that the antagonist-treated groups significantly reduced *FGFR1* expression and attenuated Wnt/β-catenin signaling, as evidence by reductions in *Wnt10b* and *Axin2*. Reduction in expression of osteoblast message levels for Type 1 *collagen*, *ALP*, and *Dmp1* were observed (**Table 3**). In contrast, mature Ob markers such as *MEPE* and *Osteocalcin* were significantly upregulated in femurs from the antagonists treated groups. In addition, adipocyte markers *Pparγ2*, *aP2*, and *lipoprotein lipase* (*Lpl*) were significantly downregulated in all antagonists treated groups (**Table 3**). The group treated with **13a** had greater effects in all gene expressed markers than did either ZINC13407541 or the **8n** group. However, there was no obvious changes in osteoclast markers including *OPG*, *RankL*, *Mmp9*, and *Trap* as well as *Fgf23* and *Fgf2* transcripts in the treated groups as compared to vehicle treated controls (**Table 3**). Compound **13a** had greater therapeutic effects on *Hyp* mouse homologue of XLH.

**Table 3.** Gene-expression profiles in bone of *Hyp* mice treated with ZINC13407541 and its analogues.

| Gene | Vehicle | ZINC13407541 | 8n | 13a | p-value |
| --- | --- | --- | --- | --- | --- |
| **Osteoblast lineage** |  |  |  |  |  |
| *Fgfr1* | 1.00±0.35<sup>a</sup> | 0.93±0.53<sup>a</sup> | 0.94±0.37<sup>a</sup> | 0.35±0.25<sup>b</sup> | 0.0258 |
| *Fgf2* | 1.00±0.32<sup>a</sup> | 0.93±0.49<sup>a</sup> | 0.85±0.34<sup>a</sup> | 0.88±0.23<sup>a</sup> | 0.8782 |
| *Fgf23* | 1.00±0.34<sup>a</sup> | 0.93±0.50<sup>a</sup> | 0.95±0.26<sup>a</sup> | 1.00±0.26<sup>a</sup> | 0.9771 |
| *Dmp1* | 1.00±0.48<sup>a</sup> | 0.77±0.18<sup>a</sup> | 0.88±0.16<sup>a</sup> | 0.38±0.16<sup>b</sup> | 0.0002 |
| *Osteopontin* | 1.00±0.25<sup>a</sup> | 0.55±0.11<sup>b</sup> | 0.56±0.16<sup>b</sup> | 0.64±0.12<sup>b</sup> | <0.0001 |
| *Bsp* | 1.00±0.34<sup>a</sup> | 0.87±0.27<sup>a</sup> | 0.95±0.29<sup>a</sup> | 0.90±0.18<sup>a</sup> | 0.8607 |
| *Mepe* | 1.00±0.16<sup>a</sup> | 1.68±0.35<sup>b</sup> | 1.86±0.44<sup>b</sup> | 5.42±0.77<sup>c</sup> | <0.0001 |
| *Col1* | 1.00±0.28<sup>a</sup> | 0.55±0.16<sup>b</sup> | 0.62±0.23<sup>b</sup> | 0.30±0.10<sup>c</sup> | <0.0001 |
| *Alp* | 1.00±0.26<sup>a</sup> | 0.66±0.20<sup>b</sup> | 0.60±0.19<sup>b</sup> | 0.36±0.08<sup>c</sup> | 0.0001 |
| *Osteocalcin* | 1.00±0.27<sup>a</sup> | 1.06±0.32<sup>a</sup> | 1.25±0.39<sup>a</sup> | 3.36±0.97<sup>b</sup> | <0.0001 |
| *Runx2-II* | 1.00±0.23<sup>a</sup> | 0.82±0.41<sup>a</sup> | 0.91±0.31<sup>a</sup> | 0.21±0.11<sup>b</sup> | 0.0003 |
| *Opg* | 1.00±0.20<sup>a</sup> | 1.02±0.33<sup>a</sup> | 0.96±0.29<sup>a</sup> | 0.92±0.37<sup>a</sup> | 0.9488 |
| *RankL* | 1.00±0.16<sup>a</sup> | 0.85±0.18<sup>a</sup> | 0.97±0.37<sup>a</sup> | 0.87±0.11<sup>a</sup> | 0.6112 |
| *Fzd2* | 1.00±0.38<sup>a</sup> | 0.51±0.21<sup>b</sup> | 0.62±0.16<sup>b</sup> | 0.17±0.08<sup>c</sup> | <0.0001 |
| *Wnt10b* | 1.00±0.27<sup>a</sup> | 0.53±0.24<sup>b</sup> | 0.64±0.16<sup>b</sup> | 0.12±0.06<sup>c</sup> | <0.0001 |
| *Axin2* | 1.00±0.25<sup>a</sup> | 0.57±0.21<sup>b</sup> | 0.67±0.17<sup>b</sup> | 0.11±0.06<sup>c</sup> | <0.0001 |
| **Osteoclast** |  |  |  |  |  |
| *Trap* | 1.00±0.17<sup>a</sup> | 0.91±0.24<sup>a</sup> | 1.08±0.27<sup>a</sup> | 0.86±0.22<sup>a</sup> | 0.3383 |
| *Mmp9* | 1.00±0.21<sup>a</sup> | 0.81±0.39<sup>a</sup> | 1.10±0.34<sup>a</sup> | 1.11±0.13<sup>a</sup> | 0.2358 |
| **Chondrocyte** |  |  |  |  |  |
| *Collagen II* | 1.00±0.45<sup>a</sup> | 3.72±1.08<sup>b</sup> | 3.43±2.27<sup>b</sup> | 14.4±2.65<sup>c</sup> | <0.0001 |
| *VegfA* | 1.00±0.38<sup>a</sup> | 0.83±0.51<sup>a</sup> | 0.77±0.48<sup>a</sup> | 0.81±0.27<sup>a</sup> | 0.7852 |
| **Adipocyte** |  |  |  |  |  |
| *PPARγ2* | 1.00±0.32<sup>a</sup> | 0.43±0.29<sup>b</sup> | 0.48±0.31<sup>b</sup> | 0.10±0.08<sup>c</sup> | 0.0001 |
| *aP2* | 1.00±0.13<sup>a</sup> | 0.78±0.15<sup>b</sup> | 0.70±0.16<sup>b</sup> | 0.23±0.11<sup>c</sup> | <0.0001 |
| *Lpl* | 1.00±0.34<sup>a</sup> | 0.47±0.23<sup>b</sup> | 0.54±0.24<sup>b</sup> | 0.17±0.07<sup>c</sup> | <0.0001 |

Data are mean ±S.D. from 5-6 tibias of vehicle, ZINC13407541, or its analogues treated *Hyp* mice and expressed as the fold changes relative to the housekeeping gene *β-actin* subsequently normalized to control mice. Values with the different superscript are significantly different at *P*<0.05.

### Renal FGF23 signaling changes in kidney of *Hyp* mice treated long-term with FGF23 inhibitors

Four weeks of treatment with FGF23 inhibitors significantly altered FGF23 responsive gene expression in kidney of *Hyp* mice (**Table 4**). Compared with ZINC13407541 and **8n** groups, **13a** exhibited greater effects to increase *Npt2a* and *Npt2c message* expression, consistent with higher serum phosphate levels in the **13a** treated groups. Consistent with FGF23 stimulation of *Cyp27b1*, and inhibition of *Cyp24a1*, leading to reductions in circulating 1,25(OH)2D, the administration of ZINC13407541 and its analogues to *Hyp* mice increased the serum concentration of 1,25(OH)2D (**Table 2**), in association with increased *Cyp27b1* and decreased *Cy24a1* message expression (**Table 4**). FGF23 also stimulates *NCC* expression in the distal renal tubule cells leading to increased Na^+^ and K^+^ channel activity, sodium retention, and suppression of serum aldosterone. ZINC13407541, **8n**, and **13a** treatment suppressed NCC expression (**Table 4**), in association with increased circulating aldosterone levels in *Hyp* mice. In contrast, treatment with ZINC13407541, **8n**, and **13a** had no effects on *Fgfr1* expression, but significantly increased *Klotho* transcripts in kidney of *Hyp* mice (**Table 2**). These data indicate that ZINC13407541 and its analogues may have clinical utility in blocking the effects on the kidney of excess FGF23.

**Table 4.**
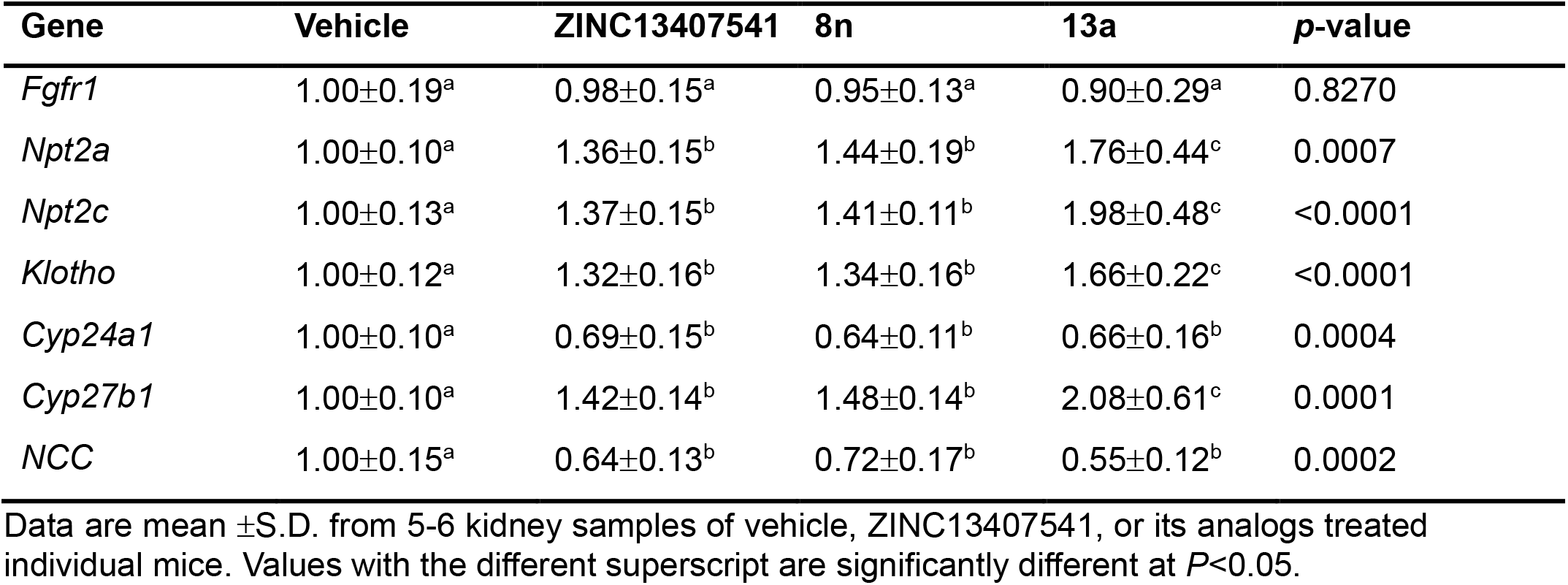
Gene-expression profiles in kidney of *Hyp* mice treated with ZINC13407541 & its analogues.

### Potential off-target effects of compound 8n

In contrast to the specificity of blocking antibodies, small molecules have a greater potential for off-target effects. Indeed, radioligand binding assays revealed that compound **8n** has greater than 50% inhibition of androgen, serotonin, dopamine, and norepinephrine receptors and activation of histamine receptor (**Table 5 and Table 1S**).

**Table 5.** Potential off-target effects of compound 8n in the radioligand binding assays.

| Assay Name | Species | Concentration |  | % inhibition<br>or stimulation |
| --- | --- | --- | --- | --- |
| Androgen (Testosterone) | Human | 10 μM | | 79 |
| Histamine H2 | Human | 10 μM | | -54 |
| Serotonin 5-HT2B | Human | 10 μM | | 63 |
| Transporter, Dopamine (DAT) | Human | 10 μM | | 86 |
| Transporter, Norepinephrine (NET) | Human | 10 μM | | 75 |

## DISCUSSION

We are pursuing the development of small molecules that bind to FGF23 and inhibit its activation of the FGFR/α-Klotho binary receptor complex. Herein we report on the efficacy of ZINC13407541 derivatives that displayed either greater stability in *in vitro* microsomal assays or greater potency in *in vitro* functional assays. Compound **8n** and **13a** (most potent) both demonstrated short- and long-term efficacy in blocking FGF23 end-organ effects in *Hyp* mice, a human homologue of XLH, used in pre-clinical assessment of KRN23. We found that pharmacological inhibition of FGF23 efficiently abrogates aberrant FGF23 signaling and corrects the hypophosphatemia caused by elevated FGF23 levels in *Hyp* mice. Increases in serum phosphate levels were observed as early as 3 to 5 hours after intraperitoneal injections of these compounds. The minimal effective dose was 50 mg/kg body weight.

Long-term FGF23 inhibition with **8n** and **13a** treatment of *Hyp* mice for 4 weeks resulted in sustained increases in serum phosphate and 1,25(OH)_2_D levels, enhanced bone growth, increased bone mineralization, and improvement of the rachitic growth plate abnormalities. **13a** had greater therapeutic effects on FGF23-induced abnormalities in *Hyp* mice, consistent with its lower IC_50_. The improvement in biochemistries in *Hyp* mice after treatment with these chemical FGF23 inhibitors was associated with reversal of FGF23’s renal effects. Elevated FGF23 in *Hyp* mice suppresses the phosphate transporters *Npt2a* and *Npt2c* and *Cyp27b1* the enzyme that 1-hydroxylates 25(OH)D, and suppresses *Cyp24a1*, which degrades 1,25(OH)2D in the proximal tubule. Consistent with the increase in serum phosphate and 1,25(OH)2D levels, **8n** and **13a** treatment increased *Npt2* and *Cyp27b1* expression and decreased *Cyp24a1* message expression.

Excess FGF23 in *Hyp* mice also leads to stimulation of the sodium chloride co-transporter in the distal tubule that leads to sodium retention and suppression of aldosterone ^(30)^. Treatment with **8n** and **13a** also increased aldosterone levels and decreased expression of *NCC*, the sodium chloride transporter in the distal tubule. FGF23 also suppresses α-*Klotho* message expression in the distal tubule in *Hyp* mice. Treatment with **8n** and **13a** also increased α-Klotho transcript levels in *Hyp* mice. Thus, these small molecule FGF23 inhibitors antagonized both the proximal and distal tubular effects of FGF23 excess.

Excess FGF23 has both indirect effects on bone and cartilage mineralization mediated by the reductions in serum phosphate and 1,25(OH)_2_D ^(31, 32)^, as well as possible direct effects due to FGF23 activation of FGFRs in osteoblasts ^(33, 34)^. We observed substantial improvement in bone abnormalities after only 4 weeks of treatment as well as alteration in gene expression profiles in bone. Treatment with **8n** and **13a** altered osteoblast function as evidence by decreased *Runx2-II*, *Alp*, *Col1*, *Opn* ^(35)^, and *Dmp1* and increased *Mepe*, *Bsp*, and *Osteocalcin* expressions. FGF23, may also have effects on adipocytes to modulate adipogenesis ^(30)^. This may explain the effects of FGF23 antagonists in reducing adipogenic markers in *Hyp* bone. Treatment with **8n** and **13a** significantly reduced *Fgfr1*, *Wnt10b*, *Fzd2*, and *Axin2* gene transcripts, consistent with our previous findings that *Hyp* mice exhibited higher Fgfr1 and Wnt signaling ^(27) (28)^. In addition, treatment with **8n** and **13a** did not increase the *FGF23* transcripts and serum FGF23 levels in *Hyp* mice, which differs from increased serum total FGF23 levels following treatment of XLH patients with FGF23 blocking antibodies (https://www.ultragenyx.com/file.cfm/95/docs/Crysvita_Full_Prescribing_Information.pdf)

Off-target effects of small molecule inhibitors are a potential liability compared to the high target specificity of blocking antibodies. In this regard, **8n** blocked the androgen receptor, as well as dopamine, norepinephrine, and serotonin receptors and activated the histamine receptor *in vitro*. Additional chemical modification may be needed to further increase the specificity of these leads for FGF23.

The short half-life of small molecules may be of considerable clinical benefit in titrating and reversing drug effects, which may be particularly important given the narrow therapeutic window for FGF23 suppression, particularly in chronic kidney disease (CKD). Preclinical CKD models show that inhibiting FGF23 with a blocking antibody increases mortality due to oversuppression of FGF23 ^(15)^. Drawing analogies to treatment of secondary hyperparathyroidism with short acting calcimimetics in CKD to lower PTH levels, and pointing to data that even partial reductions of FGF23 may be sufficient to reduce cardiovascular effects, a short-acting, titratable FGF23 antagonist might reduce FGF23 dose-dependent cardiotoxicity with acceptable side effects in CKD ^(1, 36, 37)^. Further studies are needed to test the effects of these small molecules in models of CKD.

In conclusion, these studies further advance the premise that the development of FGF23 small molecule inhibitors is a novel approach for treatment of FGF23-mediated hypophosphatemic diseases. Small molecules are generally more cost-effective, have a longer shelf life, and are more easily distributed. In addition, an oral anti-FGF23 therapy with small molecule inhibitors has the potential to expand the current treatment repertoire by increasing patient satisfaction and adherence.

## MATERIALS AND METHODS

### Molecular docking

To computationally analyze the binding conformation and affinity of ZINC13407541 and its two analogs, **13a** and **8n**, to FGF23, molecular docking of the three compounds on FGF23 was performed using AutoDock VinaMPI ^(21, 22)^. The crystal structure of FGF23 [PDB code: 5W21^(38)^] was used as the receptor, and a pose-searching box of 40 x 30 x 40 Å was centered at the geometric center of Gly38 to Arg161 to include the whole N-terminal domain of FGF23. An exhaustiveness of 30 was used for adequate sampling of ligand conformations within the box. Following docking, the pose with the best score (i.e., the lowest value indicating estimated free energy of binding, ΔG) for each compound was plotted in 3D and 2D using VMD ^(39)^ and LigPlot^+ (40)^, respectively. For hydrogen-bond calculation in LigPlot^+^, the maximum hydrogen-acceptor and maximum donor-acceptor distances were set at 3.0 and 4.0 Å, respectively. For non-bonded contact calculation, the minimum and maximum contact distances were set at 2.0 and 4.0 Å, respectively. K_DEEP_ ^(23)^, a protein-ligand binding affinity predictor based on deep convolutional neural networks, was also used to obtain ΔG for each compound as a comparison with AutoDock VinaMPI ^(21, 22)^.

### Chemicals and reagents

Synthetic preparation of new ZINC13407541 analogues was conductend in the medicinal chemistry lab at UTHSC College of Pharmacy and Tennessee Technological University. All reagents used for organic synthesis were sourced from Sigma-Aldrich (St. Louis, MO) and Fisher Scientific (Fairfield, NJ), and stored in accordance with manufacturer recommendations. Each test compound was fully characterized by mass spectrometry and NMR (^1^H and ^13^C), with a >95% chemical purity which was determined with HPLC. The medicinal chemistry group also generated gram-scale quantitates for *in vivo* animal studies as we previously reported ^(24)^. The compounds were stored at −20 °C freezer and were tested both *in vitro* assays and *in vivo* in the laboratories.

Recombinant human FGF23 and FGF2 were purchased from R&D Systems (Minneapolis, MN, USA). Synthetic human C-terminal FGF23 (residues 180-251) peptide (FGF23CT), which binds to α-KL and blocks full-length FGF23 binding to the FGFR/α-KL binary complex, was obtained from Phoenix Pharmaceuticals, Inc. (Burlingame, CA, USA). Recombinant human FGF-, FGF-19, and FGF-21 were purchased from PeproTech (Rocky Hill, NJ, USA).

### Cell culture and in vitro functional assays

HEK293T cells were cultured in DMEM containing 10% FBS and 1% penicillin and streptomycin (P/S).To test the effects of the novel compounds on FGF23-mediated activation of FGFR1/α-KL complex, HEK293T cells were transiently transfected with either empty expression vector or full-length human α-KL along with the ERK luciferase reporter system ^(41)^ and *Renilla* luciferase-null as internal control plasmid. Transfections were performed by electroporation using Cell Line Nucleofector Kit R according to the manufacturer’s protocol (Amaxa, Inc., Gaithersburg, MD). Thirty-six hours after transfection, the transfected cells were treated with the test compound with a range of 10^−9^~10^−4^ M in the presence or absence of 1 nM FGF23. After 5 hours, the cells were lysed and luciferase activities measured using a Synergy H4 Hybrid Multi-Mode Microplate Reader (Winooski, VT, USA) and Promega Dual-Luciferase Reporter Assay System (Madison, WI, USA).

### *In vitro* ADME and *in vivo* pharmacokinetics (PK) studies

We contracted the *in vitro* ADME screening and *in vivo* PK studies with Eurofins Pharma Discovery Services. Microsomal stability of ZINC13407541 and ZINC13407541 analogues was determined in liver microsomes, including human liver microsomes (HLM) and mouse liver microsomes (MLM) to determine potential species differences in metabolism ^(42)^. Aqueous solubility was assessed in PBS (pH 7.4), simulated intestinal fluid, and simulated gastric fluid. We also extended our assessment of compound *in vitro* ADME by assessing protein binding, permeability, and metabolism. We assessed protein binding using rapid equilibrium dialysis, as previously described ^(43)^. A study on single dose pharmacokinetics after intravenous (i.v.) and oral administration determined basic PK parameters including clearance, volume of distribution, elimination half-life, systemic exposure, residence time and oral bioavailability. Mice (24/route of administration) received a single oral or i.v. (1~10 mg/kg) dose of nominated analogues. Mice (n=3/time point) were sacrificed at 0, 5, 15, 30, 60, 90, 120, 240, 360 and 1440 min. Plasma were stored at −80°C until analysis by LC-MS/MS. PK parameters (bioavailability, clearance, and half-life) were determined using non-compartmental methods ^(44)^.

### RNA purification and quantitative real-time RT-PCR

For quantitative real-time RT-PCR, 1.0 μg total RNA isolated from heart, kidney or long bone of 8-week-old mice was reverse transcribed as previously described ^(45, 46)^. PCR reactions contained 100 ηg template (cDNA or RNA), 300 ηM each forward and reverse primers, and 1XqPCR Supermix (Bio-Rad, Hercules, CA) in 50 μl. The threshold cycle (Ct) of tested-gene product from the indicated genotype was normalized to the Ct for cyclophilin A. Expression of total *Klotho* isoform transcripts was performed using the following *Klotho-*isoform-specific primers: Forward primer of mouse *m-KL*^135^ transcript: 5’-CAT TTC CCT GTG ACT TTG CTT G-3’, and reverse primer: 5’-ATG CAC ATC CCA CAG ATA GAC-3’. Forward primer of mouse *s-KL^70^* transcript: 5’-GAG TCA GGA CAA GGA GCT GT-3’, and reverse primer: 5’-GGC CGA CAC TGG GTT TTG-3’. The sequences of primers for kidney and bone gene transcripts were previously reported ^(45, 46)^. In addition, the fold change of tested gene transcripts was calculated from the relative levels of the normal gene transcripts in wild-type mice.

### Specificity binding assays by radioligand binding assays of small molecule 8n

We contracted the specificity binding assays of compound **8n** with Eurofins Panlabs Discovery Services Taiwan, Ltd. Radioligand binding assays were performed using LeadProfilingScreen (Total number of Assays: 67) and compound **8n** at a concentration of 10 μM. Biochemical assay results presented as the percent inhibition of specific binding or activity throughout the report. Significant responses (≥ 50% inhibition or stimulation for biochemical assays) were noted in the primary assays listed ^(47, 48)^.

### Animal experiments

All animal research was conducted according to guidelines provided by the National Institutes of Health and the Institute of Laboratory Animal Resources, National Research Council. The University of Tennessee Health Science Center’s Animal Care and Use Committee approved all animal studies (Protocol number: 18-111.0). All mice were maintained in our vivarium on a standard diet (7912; Harlan Teklad, Madison, WI, USA). To generate hemizygous *Hyp* mice, we crossed male hemizygous with female *wild-type* to obtain both male and female hemizygous *Hyp* mice as previously described ^(28)^. At 4 weeks of age, hemizygous *Hyp* mice were randomly assigned to the following experiments. For single-dose experiment, the *Hyp* mice received a single IP dose of ZINC13407541 (0, 50, 100, 200 mg/kg) or **13a** (0, 10, 50, 100 mg/kg) at time 0 and were collected serum samples 0, 2, 4, 8, and 24 hours after administration. For short-term efficacy studies, the *Hyp* mice were divided into four different groups: (1) Vehicle control; (2) ZINC13407541 (100 mg/kg); (3) **8n** (100 mg/kg); and (4) **13a** (100 mg/kg). The mice were treated with intraperitoneal injection of either vehicle control (5% DMSO in corn oil), or ZINC13407541, or **8n**, or **13a** twice a day for 3 days. For long-term treatments, the *Hyp* mice were divided into four different groups: (1) Vehicle control; (2) ZINC13407541 (50 mg/kg); (3) **8n** (50 mg/kg); and (4) **13a** (50 mg/kg). The mice were treated with intraperitoneal injection of either vehicle control (5% DMSO in corn oil), or ZINC13407541, or **8n**, or **13a** twice a day for four weeks. We assessed the effects of the compounds on the skeletal phenotype at 8 weeks of age (after 4 weeks treatment) using the methods previously described in studies that characterize the *in vivo* phenotypes of *Hyp* mice ^(28)^. The serum samples were collected 4 hours after last dose administration. Serum FGF23 levels were measured using the FGF23 ELISA kit (Kainos Laboratories, Tokyo, Japan). Serum phosphorus levels were measured using a Phosphorus Liqui-UV kit (Stanbio Laboratories, Boerne, TX, USA) and serum calcium levels were measured using a Calcium (CPC) Liquicolor kit (Stanbio Laboratories, Boerne, TX, USA). Serum PTH levels were measured using the Mouse Intact PTH ELISA kit (Immutopics, Carlsbad, CA, USA). Serum aldosterone levels were measured using aldosterone ELISA Kit (Cayman Chemical, MI, USA). Serum 1,25(OH)_2_D levels were measured using the 1,25-dihydroxy-vitamin D EIA Kit (Immunodiagnostic Systems, Fountain Hills, AZ, USA) as previously escribed ^(28)^.

### Bone densitometry, histomorphometric, and micro-CT analysis

Bone mineral density (BMD) of femurs was assessed at 8 weeks of age using a small animal bone densitometer (Lunar Corp, Madison, WI). Calcein (Sigma-Aldrich, St. Louise, MO) double labeling of bone and histomorphometric analyses of periosteal mineral apposition rate (MAR) in tibias were performed using the osteomeasure analysis system (Osteometrics, Inc., Decatur, GA). The distal femoral metaphyses were also scanned using a micro-CT 40 scanner (Scanco Medical AG, Brüttisellen, Switzerland). A 3D images analysis was done to determine width of growth plate (GP), bone volume (BV/TV) and cortical thickness (Ct.Th) as previously described ^(49, 50)^.

### Statistical analysis

We evaluated differences between two groups by unpaired t-test, multiple groups by one-way analysis of variance, and two groups over time by two-way analysis of variance with interactions (Table S3). All values are expressed as means ± S.D. All computations were performed using a commercial biostatistics software (GraphPad Software Inc. La Jolla, CA).

## Acknowledgement

This work was supported by grant R01-DK121132 to LDQ from the National Institutes of Health. K.L. and L.P. were sponsored by the Laboratory Directed Research and Development Program of Oak Ridge National Laboratory. This research also used resources of the Compute and Data Environment for Science (CADES) at the Oak Ridge National Laboratory, which is supported by the Office of Science of the U.S. Department of Energy under Contract No. DE-AC05-00OR22725.

