## Supplemental Figure 1S and Table 1S for "Small molecule FGF23 inhibitors increase serum phosphate and improve skeletal abnormalities in *Hyp* mice"

**Supplemental Data**

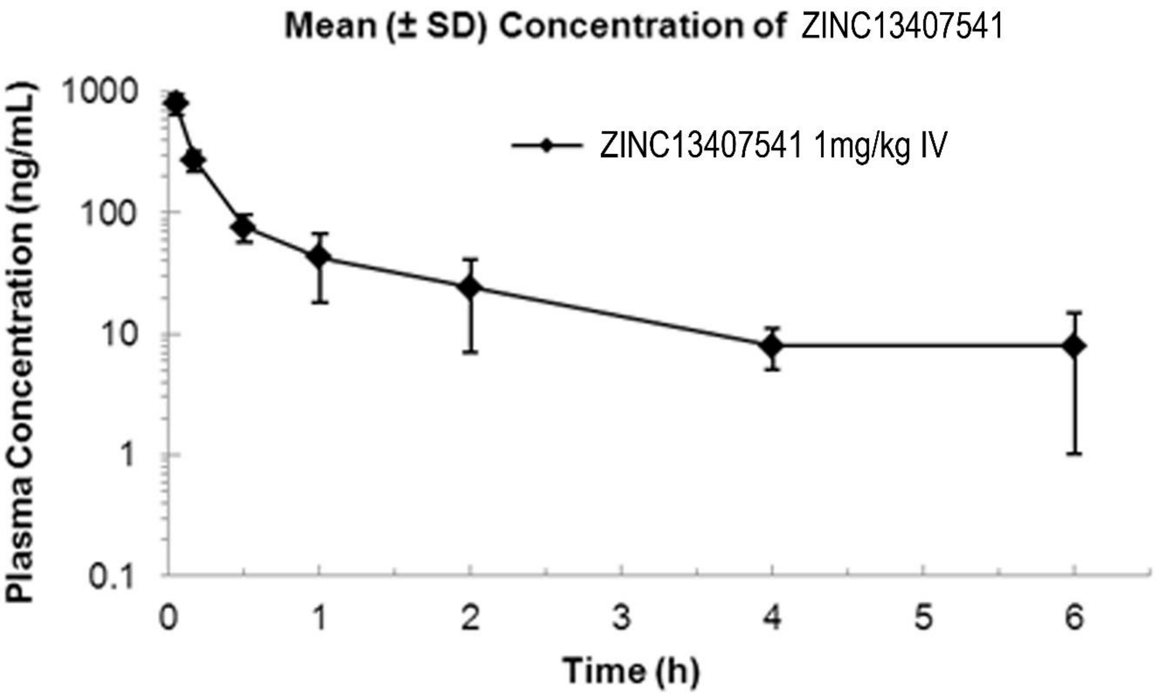


**Figure 1S.** ZINC13407541pharmacokinetics in mice (n=3) following IV administration of a single dose (1mg/kg). Data represent the mean ± SD.

**Table 1S. Potential off-target effects of compound 8n in the radioligand binding assays.**

| Assay Name | Species | Concentration | % inhibition or stimulation |
|---|---|---|---|
| Adenosine $A_1$ | Human | 10 µM | 33 |
| Adenosine $A_{2A}$ | Human | 10 µM | 16 |
| Adenosine $A_3$ | Human | 10 µM | 45 |
| Adrenergic $\alpha_{1A}$ | Human | 10 µM | 5 |
| Adrenergic $\alpha_{1B}$ | Human | 10 µM | -15 |
| Adrenergic $\alpha_{1D}$ | Human | 10 µM | 7 |
| Adrenergic $\alpha_{2A}$ | Human | 10 µM | 14 |
| Adrenergic $\beta_1$ | Human | 10 µM | 4 |
| Adrenergic $\beta_2$ | Human | 10 µM | 12 |
| Androgen (Testosterone) | Human | 10 µM | 79 |
| Bradykinin $B_1$ | Human | 10 µM | -3 |
| Bradykinin $B_2$ | Human | 10 µM | -27 |
| Calcium Channel L-Type, Benzothiazepine | rat | 10 µM | 21 |
| Calcium Channel L-Type, Dihydropyridine | rat | 10 µM | 49 |
| Calcium Channel N-Type | rat | 10 µM | -6 |
| Cannabinoid $CB_1$ | Human | 10 µM | 20 |
| Dopamine $D_1$ | Human | 10 µM | 30 |
| Dopamine $D_{2S}$ | Human | 10 µM | 12 |
| Dopamine $D_3$ | Human | 10 µM | 23 |
| Dopamine $D_{4.4}$ | Human | 10 µM | -15 |
| Endothelin $ET_A$ | Human | 10 µM | 11 |
| Endothelin $ET_B$ | Human | 10 µM | 0 |
| Epidermal Growth Factor (EGF) | Human | 10 µM | 2 |
| Estrogen ERα | Human | 10 µM | 6 |
| $GABA_A$, Flunitrazepam, Central | rat | 10 µM | -5 |
| $GABA_A$, Muscimol, Central | rat | 10 µM | 16 |
| Glucocorticoid | Human | 10 µM | -4 |
| Glutamate, Kainate | rat | 10 µM | 9 |
| Glutamate, NMDA, Agonism | rat | 10 µM | 5 |
| Glutamate, NMDA, Glycine | rat | 10 µM | -2 |
| Glutamate, NMDA, Phencyclidine | rat | 10 µM | 0 |
| Histamine $H_1$ | Human | 10 µM | 42 |
| Histamine $H_2$ | Human | 10 µM | -54 |

**Table 1S (continued).**

| | | | |
|---|---|---|---|
| Histamine $H_3$ | Human | 10 µM | 0 |
| midazoline $I_2$, Central | rat | 10 µM | 3 |
| Interleukin IL-1 R1 | Human | 10 µM | 19 |
| Leukotriene, Cysteinyl $CysLT_1$ | Human | 10 µM | 25 |
| Melatonin $MT_1$ | Human | 10 µM | -3 |
| Muscarinic $M_1$ | Human | 10 µM | 0 |
| Muscarinic $M_2$ | Human | 10 µM | 7 |
| Muscarinic $M_3$ | Human | 10 µM | 9 |
| Neuropeptide Y $Y_1$ | Human | 10 µM | -6 |
| Neuropeptide Y $Y_2$ | Human | 10 µM | -11 |
| Nicotinic Acetylcholine α1, Bungarotoxin | Human | 10 µM | 0 |
| Nicotinic Acetylcholine α3β4 | Human | 10 µM | -1 |
| Opiate $\delta_1$ (OP1, DOP) | Human | 10 µM | 7 |
| Opiate κ (OP2, KOP) | Human | 10 µM | 0 |
| Opiate µ (OP3, MOP) | Human | 10 µM | 9 |
| Phorbol Ester | mouse | 10 µM | 6 |
| Platelet Activating Factor (PAF) | Human | 10 µM | -14 |
| Potassium Channel [KATP] | Hamster | 10 µM | -11 |
| Potassium Channel hERG | Human | 10 µM | 25 |
| Prostanoid $EP_4$ | Human | 10 µM | 34 |
| Purinergic P2X | rat | 10 µM | -13 |
| Purinergic P2Y, Non-Selective | rat | 10 µM | 2 |
| Rolipram | rat | 10 µM | -2 |
| Serotonin (5-Hydroxytryptamine) 5-$HT_{1A}$ | Human | 10 µM | -42 |
| Serotonin (5-Hydroxytryptamine) 5-$HT_{2B}$ | Human | 10 µM | 63 |
| Serotonin (5-Hydroxytryptamine) 5-$HT_3$ | Human | 10 µM | -5 |
| Sigma σ1 | Human | 10 µM | 33 |
| Sodium Channel, Site 2 | rat | 10 µM | 37 |
| Tachykinin $NK_1$ | Human | 10 µM | -2 |
| Thyroid Hormone | rat | 10 µM | 16 |
| Transporter, Dopamine (DAT) | Human | 10 µM | 86 |
| Transporter, GABA | rat | 10 µM | 4 |
| Transporter, Norepinephrine (NET) | Human | 10 µM | 75 |
| Transporter, Serotonin (5- Hydroxytryptamine) (SERT) | Human | 10 µM | 13 |
